# *Naegleria* amoebae seek confinement and crawl persistently through narrow spaces

**DOI:** 10.1101/2025.11.30.691445

**Authors:** Katrina B. Velle, Meera Ramaswamy, Babak Vajdi Hokmabad, Tania Martín-Pérez, Teodoro Tapia Carrasco, William S. Callahan, Harrison S. Kim, Emily M. Larkin, Abdurrahman H. ElZafarany, Samantha M. Jacques, Sujit S. Datta, Marc Edwards, Lillian K. Fritz-Laylin

## Abstract

The "brain-eating amoeba" *Naegleria fowleri* dwells in ponds where it normally feeds on bacteria, but if it enters the brain it can cause a deadly infection. To establish infection, *N. fowleri* must migrate through different environments—along olfactory axons, through openings in the cribriform plate, and within brain tissue—yet how it does so remains unknown. As a model for *N. fowleri* migration within these environments, we examine how its non-pathogenic relative, *Naegleria gruberi*, navigates environments of distinct geometries. We show that *Naegleria* uses both actin-rich protrusions and membrane blebs to crawl across or between flat surfaces. We also explore how *Naegleria* interact with narrow channels and find that, unlike *Dictyostelium* amoebae that we show frequently disengage from channel interfaces, *Naegleria* amoebae probe channels until they enter. Once inside, *Naegleria* crawls quickly (>50 μm/min) and unidirectionally over long distances (>1 mm) using only bleb-based motility. We also introduced *Naegleria* to granular hydrogel matrices that mimic pond sediments and found that cells readily enter and migrate through these three-dimensional matrices using both blebs and lamellar protrusions. Although cells in matrices showed lower persistence at short timescales, longer time scales correlate with increased persistence, suggesting *Naegleria* cells may retain memory of past orientation. We propose that pond life may select for three behaviors that prime *Naegleria* for pathogenesis: memory-guided motility that would facilitate exploration of sinus cavities, confinement-seeking (“claustrophilia”) that would promote entry into narrow passages along olfactory axons, and persistent bleb-based migration that would allow rapid transit along axons to the brain.

## INTRODUCTION

Many human pathogens are opportunistic: under normal conditions, they do not cause disease, but in a particular context, they become lethal. Understanding which features of these normally benign organisms enable pathogenesis is central to understanding the diseases they cause. For example, the “brain-eating amoeba” *Naegleria fowleri* is a pond-dwelling protist that normally feeds on bacteria (Stahl and Olson 2020; Fulton 1970). If it gains access to the brain, however, it can cause a devastating infection known as primary amoebic meningoencephalitis (Siddiqui et al. 2016; Moseman 2020). Sadly, most *Naegleria* infections occur in children, and the lack of reliable treatments has led to an astronomical case fatality rate of ∼95% (Hall et al. 2024). Despite its lethality, little is known about the basic biology of *N. fowleri—*or any *Naegleria* species, even the non-pathogenic model *Naegleria gruberi.* This lack of information makes it difficult to understand which aspects of its biology prime *Naegleria* for infection or how those features function in its normal environment.

While the mechanisms underlying the ability of *N. fowleri—*but not other *Naegleria* species*—*to infect humans remain mysterious (Liechti et al. 2018; Herman et al. 2021), what is clear is that *Naegleria* must move from its native environment into a human brain to establish infection. *N. fowleri* infections typically begin when water forcibly enters the nose, often during freshwater recreation (Hall et al. 2024). Once in the upper sinuses, *N. fowleri* crawls along olfactory axons (Jarolim et al. 2000), which form tight bundles that pass through small openings in the cribriform plate—a piece of bone that separates the brain from the sinuses. To penetrate the cribriform plate and spread through brain tissue, *N. fowleri* must crawl through confined spaces. Although crawling in confinement is essential for *N. fowleri* pathogenesis, this behavior remains almost entirely unstudied.

Many cells, including white blood cells and *Dictyostelium* amoebae, use distinct mechanisms when crawling across flat surfaces versus in confinement (Fritz-Laylin and Titus 2023). When crawling across flat surfaces, these cells form lamellar protrusions that are filled with branched-actin networks that push the membrane forward (Mullins et al. 1998; Bieling and Rottner 2023). Confinement induces animal cells to switch to an alternative form of crawling: blebbing motility, which is driven by the eruptive detachment of the plasma membrane from the underlying actin cortex (Liu et al. 2015; Graziano et al. 2019; García-Arcos et al. 2024). A similar confinement-induced switch from lamellar protrusions to membrane blebs has been demonstrated in *Dictyostelium* amoebae (Srivastava et al. 2020; Zatulovskiy et al. 2014). Moreover, choanoflagellates, which are unicellular relatives of animals that usually swim, can also bleb when confined—a response thought to help escape entrapment in silt (Brunet et al. 2021). Our prior work shows that *Naegleria gruberi* uses lamellar protrusions when crawling on coverslips (Velle and Fritz-Laylin 2020). The suggestion that *Naegleria* may also form blebs (Russell et al. 2017), along with the widespread distribution of confinement-induced blebbing among diverse eukaryotic cell types, raises the possibility that *Naegleria* may similarly use blebbing in confined environments, including those they encounter during infection.

To explore how *Naegleria* might crawl in normal habitats and in disease contexts, we examined *N. gruberi’s* migration in four distinct physical environments: across flat surfaces, sandwiched between flat surfaces, squeezed within microchannels, and through complex, three-dimensional granular matrices. We show that, similar to animal and *Dictyostelium* cells (Liu et al. 2015; Graziano et al. 2019; Srivastava et al. 2020; Zatulovskiy et al. 2014), *Naegleria* switches to blebbing as the primary mode of motility when tightly confined. These confined cells crawl faster and more directionally than unconfined cells. Moreover, *Naegleria* amoebae readily enter microchannels and three dimensional matrices in the absence of chemical cues. We suggest that *Naegleria* is naturally primed to explore the confined nooks and crannies within pond environments, which are likely to harbor abundant bacterial prey. These otherwise innocuous behaviors may become deadly in a human host, enabling rapid invasion into the brain.

## RESULTS

### *Naegleria* amoebae crawl using both blebs and lamellar protrusions

*N. gruberi* amoebae are known to crawl using lamellar protrusions (Velle and Fritz-Laylin 2020) but have also been suggested to use blebs (Velle and Fritz-Laylin 2020; Russell et al. 2017). To address how frequently *Naegleria* uses each type of protrusion in different environments, we first confirmed that we could accurately differentiate actin-filled lamellar protrusions from actin-free blebs in cells crawling across glass coverslips. Because live imaging of *Naegleria* actin is currently impossible, we used DIC microscopy to image crawling cells, and then fixed and stained the actin cytoskeleton of these same cells to correlate protrusion shape with the underlying actin architecture (Velle et al. 2021). The leading edges of many *Naegleria* amoebae appeared wrinkled in live cells and stained intensely for actin polymers when fixed (**Fig. 1A, top, Video S1A**), consistent with lamellar protrusions (Zatulovskiy et al. 2014). In contrast, the leading edges of other cells appeared rounded and were devoid of actin polymer when fixed (**Fig. 1A, bottom, Video S1B**), consistent with blebs. Moreover, many cells with blebs also contained cytoplasmic actin “arcs”—curved bands of actin polymer a few microns behind the current edge of the plasma membrane **(Fig. 1A, arrowhead, Video S1B**). Actin arcs are hallmarks of blebbing in other systems—including *Dictyostelium*, mammalian cells, and *Entamoeba (Zatulovskiy et al. 2014; Maugis et al. 2010; Bergert et al. 2012)*— where they represent remnants of the actin cortex from previous rounds of blebbing. Taken together, these results establish that we can use protrusion shapes in living cells to reliably distinguish lamellar protrusions from blebs.

**Figure 1.**
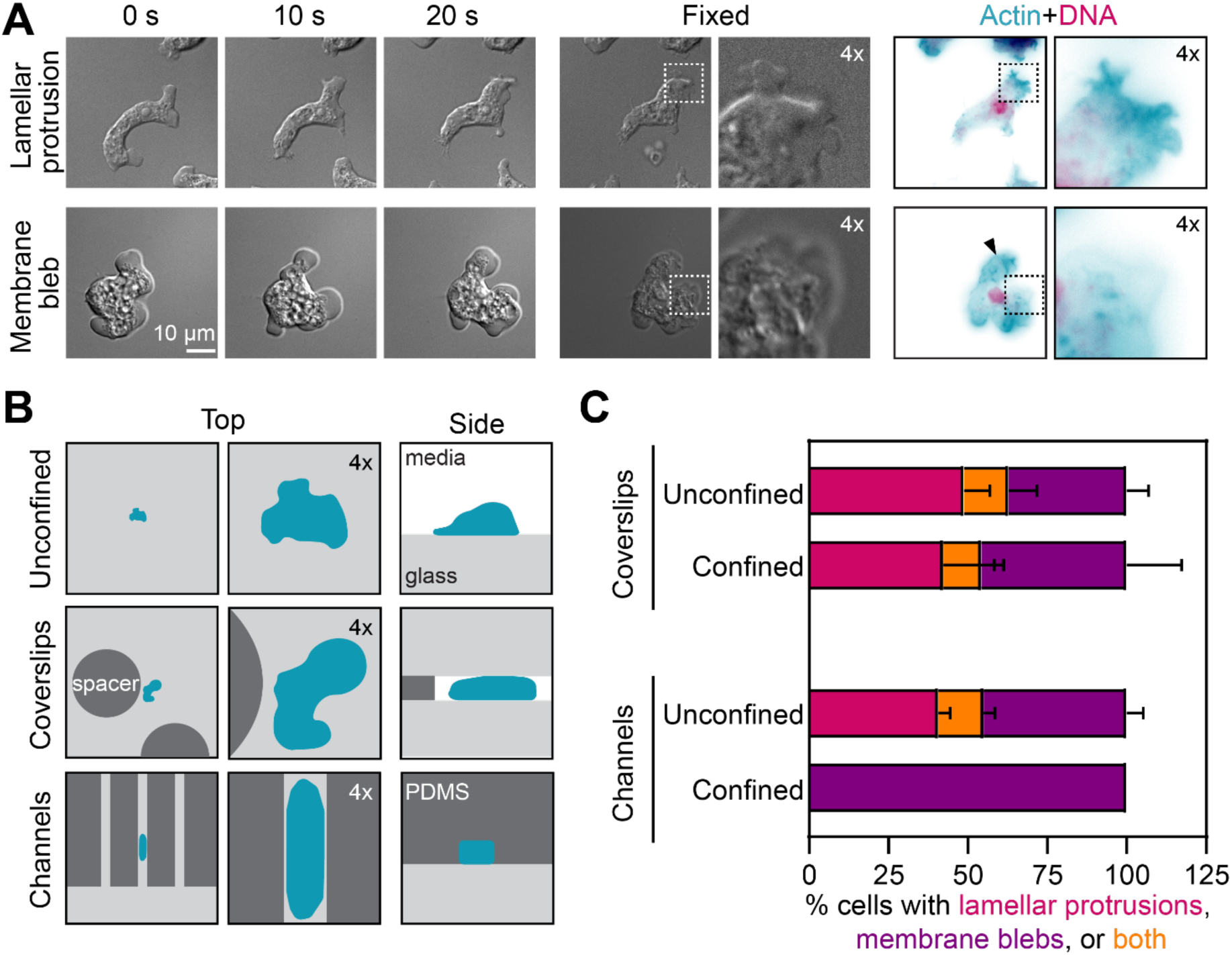
*Naegleria* uses both blebs and lamellar protrusions when crawling on or between coverslips, but relies exclusively on blebs when confined within channels. (A) *N. gruberi* cells on glass coverslips were imaged live by DIC microscopy (left panels), then fixed (middle panels) and stained *in situ* (right panels) for actin polymer using phalloidin (teal) and for DNA using DAPI (magenta). Insets show protrusions consistent with lamella (top) and blebs (bottom). **(B)** A cartoon illustrates three different environments in which amoebae (teal) were imaged. The top row shows cells crawling on glass (light gray) coverslips. The middle row shows cells crawling between glass coverslips held apart using 5 μm tall PDMS (dark gray) pillars. The bottom row shows cells crawling through 5 by 8 μm PDMS channels. **(C)** Cells imaged in the environments shown in B were scored based on protrusion types. For coverslip confinement, imaging was done before (unconfined) and after (confined) lowering the second coverslip. For channels, cells were imaged either in channels (confined) or in regions of the coverslip without channels (unconfined), and cells with obvious protrusions were categorized as having lamellar protrusions (pink), membrane blebs (purple), or both protrusion types (orange). Each bar represents the average percent of cells with each type of protrusion, with error bars indicating SD. The experiment was repeated four times on different days for confinement coverslips (with 35-128 cells scored per trial), and 3 times for channels (with 8-66 cells scored per trial).

We next wondered whether individual *Naegleria* amoebae can crawl using both lamellar protrusions and blebs. Some cells, like *Entamoeba,* zebrafish primordial germ cells, and fish keratocytes, use *either* blebs *or* lamellar protrusions to crawl (Maugis et al. 2010; Svitkina et al. 1997; Blaser et al. 2006), while others, like *Dictyostelium*, are dynamic and form both blebs and lamellar protrusions that can interconvert (Zatulovskiy et al. 2014). Tracking the protrusions of individual cells revealed that *Naegleria* amoebae exhibit a spectrum of protrusion regimes. In addition to cells with only lamellar protrusions or only blebs (**Fig. 1A**), some cells extended both blebs *and* lamellar protrusions simultaneously from different regions of the membrane (**Video S1C-D**). Moreover, while some cells formed lamellar protrusions that erupted into blebs, other cells formed blebs that subsequently extended into lamellar protrusions (**Video S1C-D**). These dynamics show that *Naegleria,* like *Dictyostelium,* can use a combination of lamellar protrusions and blebs to crawl across flat surfaces.

### When confined in microchannels, *Naegleria* crawl only by blebbing

Because other cell types that use lamellar protrusions to crawl across flat surfaces switch to blebbing motility when confined between coverslips (Liu et al. 2015; Graziano et al. 2019), we hypothesized that *Naegleria* would do likewise. To directly compare migration with and without confinement, we imaged the same cells crawling on a coverslip before and after lowering a second “confinement” coverslip held apart from the first by 5 μm spacers (Le Berre et al. 2014)(**Fig. 1B**, middle). This confinement flattened the amoebae like pancakes but did not prevent them from crawling. To quantify the protrusions produced by cells with and without confinement, we scored protrusions produced by cells that had at least one obvious protrusion before and after lowering the coverslip. Contrary to our hypothesis, the two populations appeared similar (**Fig. 1C**): comparable fractions of cells had either blebs (unconfined: 49 ± 8 %, confined: 42 ± 19 %—average ± SD), or lamellar protrusions (unconfined: 37 ± 7 %, confined: 46 ± 17 %), and fewer cells had both (confined: 14 ± 9 %, unconfined: 12 ± 4 %).

Although the *average* frequency of cells with blebs and/or lamellar protrusions were not statistically different, the marked increase in the blebbing rate in 3 of 4 individual biological replicates (**Fig. S1A**) made us wonder if we had provided sufficient confinement to trigger blebbing in some experiments but not others. This need for sufficient confinement would be consistent with work in other systems showing that high levels of confinement can be required to induce blebbing; *Dictyostelium* cells confined under agarose, for example, do not switch to blebbing unless very stiff agarose is used, or a weight is placed on top of the agarose (Srivastava et al. 2020). We therefore used a 3 μm spacer to test whether increasing confinement could boost blebbing rates. Unfortunately, this resulted in >95% of cells exploding after lowering the second coverslip (**Fig. S1B**). To increase cell confinement without bursting the cells, therefore, we turned to using narrow channels that confine cells along their sides, as well as between a top and a bottom surface (**Fig. 1B**, compare side views of cells). To determine whether and how high confinement in channels alters *Naegleria* crawling, we compared the protrusions formed by control cells outside the channels (**Fig. 1C**, bottom, “unconfined”) to those migrating within 5 μm tall by 8 μm wide channels (**Fig. 1C**, bottom, “confined”). We found that cells crawling in channels exclusively formed blebs across all three trials (100 ± 0%). To confirm that cells switch to blebbing after entering channels, we fixed and stained cells at channel entrances. Shallow protrusions that did not reach deep into channels showed intense actin staining at their tips, consistent with being lamellar. (**Fig. S2A**). Protrusions that reached deeper into channels were actin-free and contained actin arcs, consistent with blebs (**Fig. S2A-B**). These behaviors are similar to those of *Dictyostelium* cells, which transition from <50% blebbing with soft agarose overlays to nearly 100% blebbing with stiff agarose overlays (Srivastava et al. 2020). Together, these data indicate that *Naegleria* uses both lamellar protrusions and blebs when crawling across flat surfaces, but only blebs when crawling in high confinement through narrow channels. Whether and how this switch in migration impacts motility, however, was unclear.

### Speed and persistence of *Naegleria* amoebae are modulated by the degree of cell confinement

We next explored whether confinement impacts *Naegleria* crawling speed and directional persistence. We first tracked the same cohort of cells before and after lowering the 5 micron confinement coverslip (**Fig. 2A**) and found that cells crawled at 14.5 ± 4.8 μm/min prior to confinement and 20.5 ± 4.8 μm/min after (**Fig. 2B**), an increase of around 40% (p = 0.03, paired t test). To determine if this increase in speed correlates with increased blebbing, we plotted the relationship between average speed and the percent of blebbing cells. There was no obvious relationship (**Fig. S3A**), suggesting the protrusion mechanism alone may not dictate speed. We also quantified the persistence of migration—a measure of trajectory straightness—by dividing each cell’s maximum displacement from the starting point by its total path length. Using this measure, a persistence of 1 represents a cell moving in a straight line and smaller values indicate more circuitous routes. We found that the persistence of *Naegleria* amoebae is about the same with and without confinement (0.59 ± 0.03 prior to confinement, and 0.67 ± 0.06 post confinement, p = 0.07, paired t test, **Fig. 2C**). Moreover, persistence was positively correlated with speed in both unconfined and confined cells (**Fig. S3B**), similar to observations of other cell types (Maiuri et al. 2015). Taken together, these data show that confinement between coverslips primarily modulates *Naegleria* cell speed.

**Figure 2.**
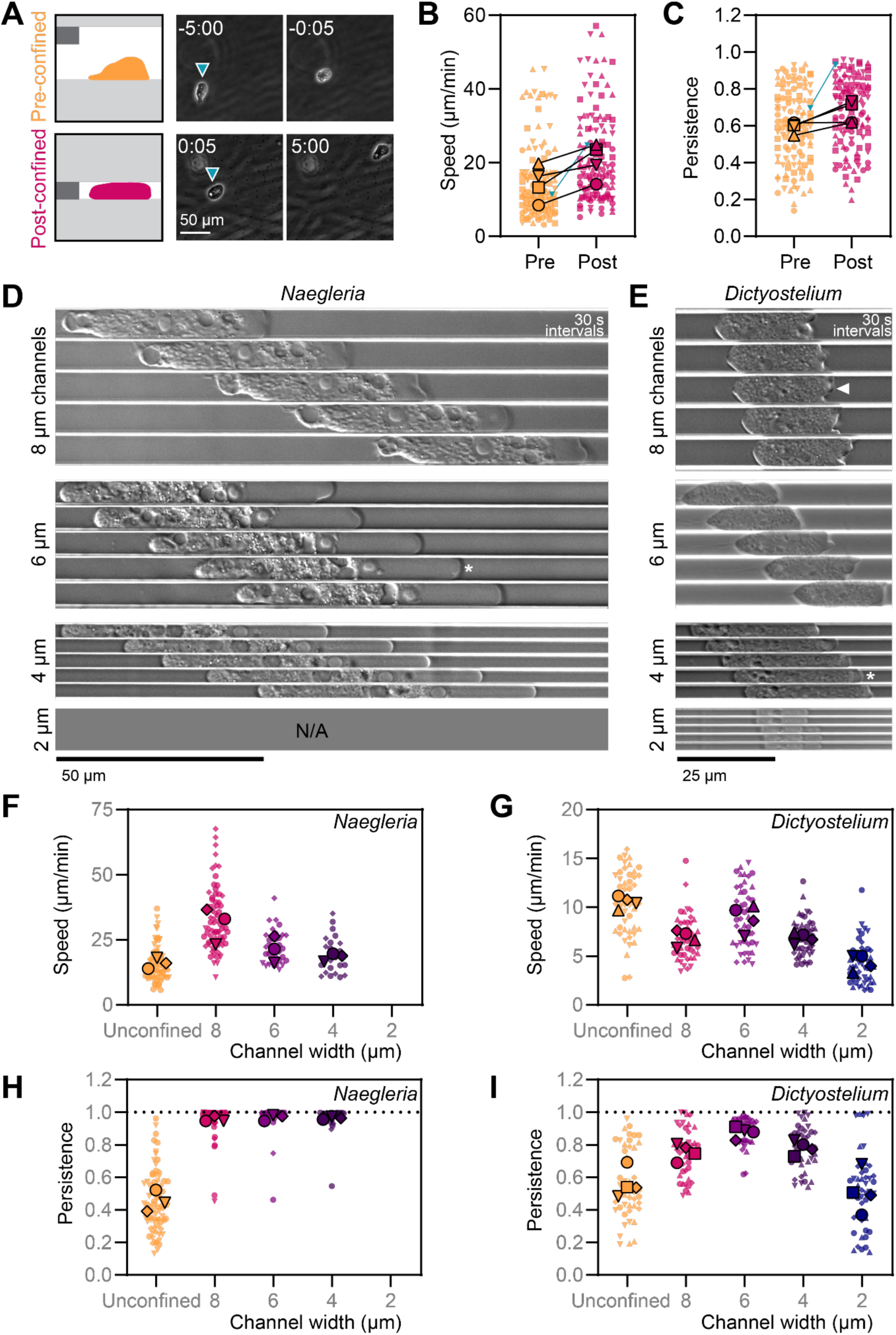
Confinement in channels correlates with fast, highly persistent movement of *Naegleria* amoebae. (A-C) *Naegleria* cells were imaged by phase contrast 20 min before and 20 min after lowering a confinement coverslip (at time 0:00 min:s) with 5 μm spacer pillars as shown in the diagram (A). Cells were tracked to measure cell speed (**B**) and maximum persistence (maximum displacement/total path length, **C**). In the SuperPlots (Lord et al. 2020) in B-C, each small symbol indicates the average value for a single cell, while large symbols show experiment-level averages, with shape coordinated by trial. The teal triangle indicates the cell shown in A. Statistics were performed on experiment level averages (B: p=0.03; C: p=0.07; paired t tests). Note: these data are from the same cells whose protrusion types are quantified in Fig. 1C. (**D-E**) *Naegleria* (**D**) and *Dictyostelium* cells (**E**) were imaged at high magnification by DIC while crawling in 2-8 μm wide channels. Representative cells are shown at 30 s intervals. Asterisks indicate examples of blebs, while an arrowhead shows a lamellar protrusion. (**F-I**) *Nageleria* and *Dictyostelium* cells were imaged in microchannels at low magnification for 45-60 min. Up to 30 *Naegleria* or *Dictyostelium* cells that fully entered the channels were tracked until they left the field of view, left the channel, or the movie ended. SuperPlots show the speeds (F, G) or persistence values (H, I) of individual cells (small symbols) and experiment-level averages (large symbols), with trials coordinated by shape. ANOVAs with Tukey’s multiple comparisons tests were performed on experiment level averages (F: unconfined vs. 8 μm, p=0.02; G: unconfined vs. 8, 4, or 2 μm, p<0.01; unconfined vs. 6 μm, p=0.11; H: unconfined vs. 8, 6, or 4 μm, p<0.01; I: unconfined vs. 8, 6, or 4 μm, p<0.02; unconfined vs. 2 μm, p=0.88). Note: *Naegleria*’s protrusion types were scored for unconfined cell vs cells confined in 8 μm channels in Fig. 1C.

We next quantified the impact of confinement in channels on *Naegleria* crawling speed and/or persistence. Because varying confinement levels alters blebbing propensity (**Fig. 1C**), we explored different degrees of confinement by using channels of different widths (8, 6, 4, and 2 μm widths, all with a 5 μm height). Although no cells entered the 2 μm wide channels, cells that entered 8, 6, or 4 μm channels again showed obvious blebbing (**Fig. 2D, Fig. S2B**). Moreover, cells confined in 8 μm channels crawled nearly twice as fast as control cells that were in the same dish but outside of channels (30.9 ± 7.0 μm/min compared to 16.0 ± 2.0 μm/min, p = 0.02 ANOVA with Tukey’s multiple comparisons test). Increasing confinement further by using 6 and 4 μm channels resulted in speeds statistically similar to unconfined cells (6 μm: 21.4 ± 5.1 μm/min, p = 0.50; 4 μm: 18.4 ± 1.5 μm/min, p = 0.91, **Fig. 2F**), indicating that moderate, but not high, confinement in channels boosts *Naegleria* crawling speed. *Naegleria* cells may move more slowly in narrower channels due to increased cell surface contact with the substrate. Plotting cell lengths versus speed of individual cells shows a negative correlation (**Fig. S3C**), and is consistent with increased surface contact leading to slower speeds. Additionally, slower movement in narrower channels could be due to squeezing of the nucleus (Le Berre et al. 2012; Stöberl et al. 2024; Wolf et al. 2013; McGregor et al. 2016), as cells in 4 micron channels show obvious nuclear deformations (**Fig. S4A**), suggesting that both mechanisms may be at play. Collectively, these data suggest that moderate confinement between coverslips *or* in microchannels induces faster crawling in *Naegleria*.

Although cells cannot bend their migration paths while crawling through straight, narrow channels, they can turn around. Because such polarity reversals lower persistence values, we next analyzed the persistence of *Naegleria* amoebae crawling through microchannels. While control cells outside the channels crawled with an average persistence of 0.44 ± 0.06, *Naegeria* amoebae became almost perfectly persistent when crawling in channels of all widths (8: 0.96 ± 0.02; 6: 0.97 ± 0.02; 4 : 0.96 ± 0.01) and rapidly moved out of the field of view (**Fig. 2H**). Unlike *Naegleria* confined between coverslips (**Fig. S3B**), we observed no clear correlation between cell speed and persistence (**Fig. S3D**). Remarkably, dozens of cells completed the entire run length of the channel, crawling nearly 2 millimeters to escape at the end (**Fig. S5**). These data demonstrate that *Naegleria* amoebae remain strongly polarized when crawling through narrow channels and almost never turn around. These experiments differ from many previously reported studies that rely on chemoattractants to lure cells into channels, a tactic that naturally biases motility in the direction of the chemoattractant (Belotti et al. 2021).

To determine whether the behavior of *Naegleria* in channels is unusual, we repeated these experiments with *Dictyostelium discoideum* amoebae. In contrast to *Naegleria, Dictyostelium* cells entered all four channel sizes, including the 2 μm channels, perhaps due to their smaller cell size, and/or their smaller nuclei (**Fig. S4B-C**). The shapes of the protrusions made by *Dictyostelium* cells in channels were consistent with both lamellar protrusions and blebs (**Fig. 2E**), unlike channel-confined *Naegleria* that only bleb. *Dictyostelium* cells also crawled significantly slower in the 2, 4, and 8 μm channels compared to unconfined control cells (p < 0.001 for all comparisons; ANOVA with Tukey’s multiple comparisons test, **Fig. 2G**). These data suggest that, unlike *Dictyostelium* amoebae which were never faster in channels than when unconfined, *Naegleria* speeds up when confined in 8 micron channels compared to unconfined migration.

We next explored whether *Dictyostelium* migrating in channels are highly polarized, similar to *Naegleria*. The persistence of *Dictyostelium* amoebae greatly increased when confined in 4-8 μm channels (p < 0.022 for all conditions, **Fig. 2I**), with an obvious peak at 6 μm. Persistence dropped to the level of unconfined cells with aggressive confinement in 2 μm channels (p = 0.877). This pattern is similar to previous work measuring *Dictyostelium* migration in channels containing chemoattractant gradients (Belotti et al. 2021), but distinct from *Naegleria*, which maintains high persistence across all channels. Furthermore, while *Dictyostelium* cells are more persistent when confined in channels >2 μm, their persistence reaches only 0.76 ± 0.05 to 0.88 ± 0.09, (**Fig. 2I**) indicating that the near-perfect persistence of *Naegleria* is not a general property of channel-confined amoebae.

### *Naegleria* enter microchannels at high rates

While imaging *Naegleria* amoebae in channels, we often observed cells accumulating at channel entrances (**Fig. 3A, right**), despite the absence of any chemoattractant or fluid flow to draw cells into the channels. Strikingly, many of these cells were extending protrusions into one or more channels (**Video S2**), hinting that these cells may be “trapped” by their propensity to enter one or more crevices (Ron et al. 2024). To explore the possibility that *Naegleria* may seek confinement, we tracked the behavior of cells at channel interfaces and categorized them as “free” when they were not in contact with the interface, in “contact” when touching the interface, “protruding” when a portion of the cell was inside one or more channels, and “inside” when the entire cell was contained within a channel (**Fig. 3**, **Fig. S6A**). We found that 65-100% of cells entered 8 μm channels across three experiments, with only 0-19% disengaging after initial contact. The remaining cells were either still contacting the interface or protruding into a channel at the end of the experiment. These behaviors changed with different channel widths: although we observed no difference in the time cells spent inserting protrusions into channels of different widths (**Fig. S6C**), fewer cells entered narrower channels, concurrent with higher rates of disengagement from the interface.

**Figure 3.**
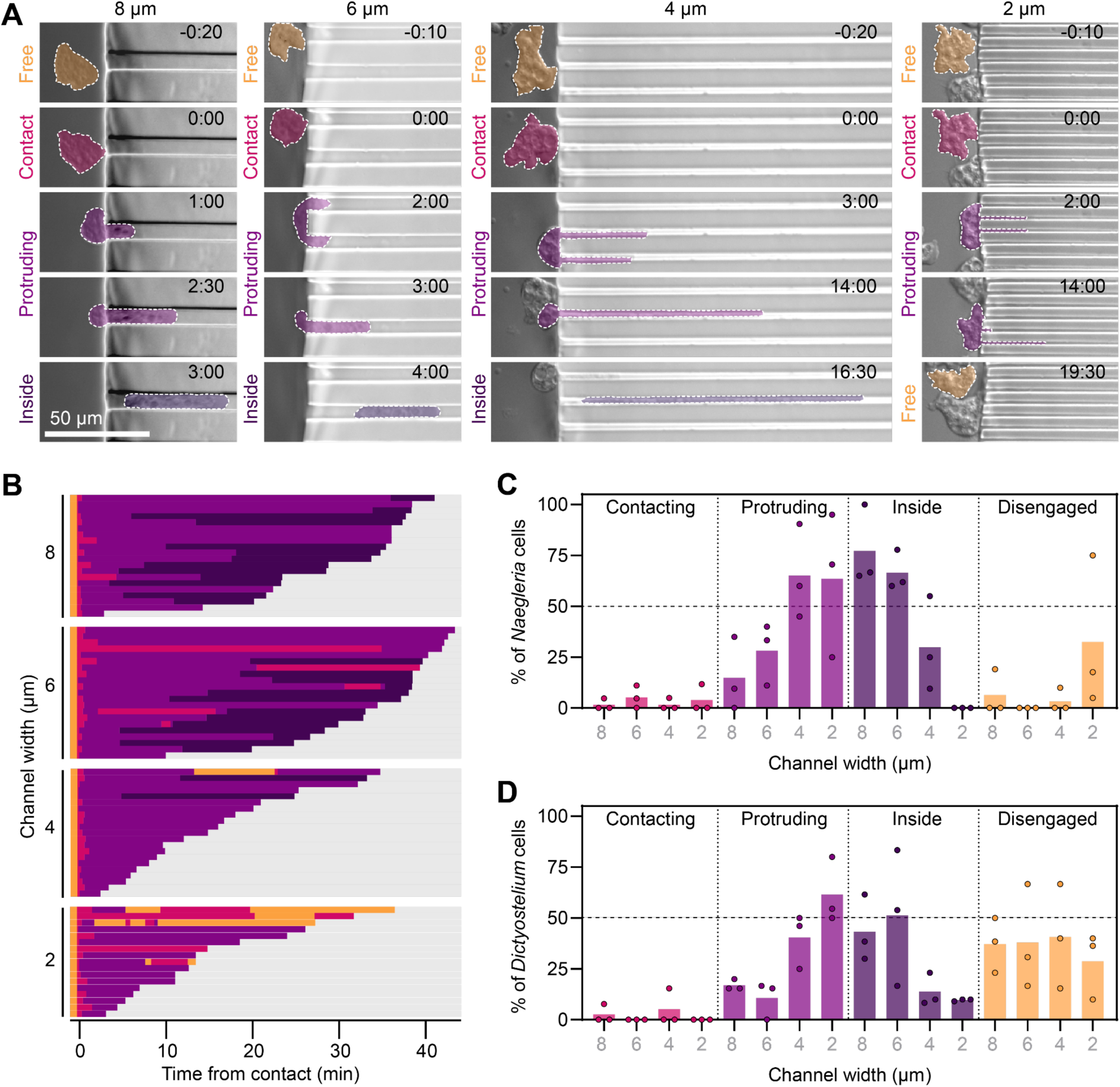
*Naegleria* cells maintain surface interactions and enter microchannels at high rates. **(A)** *Naegleria* cells that initiated interactions with channel interfaces while being imaged for experiments shown in Fig. 1C, 2F, and 2H were scored over time as being free (not in contact with the interface, orange), in contact with the interface (pink), protruding into a channel (light purple) or fully inside a channel (dark purple). **(B)** *Naegleria’s* interactions with 2-8 μm channels over 45 minutes of imaging are shown as a horizontal bar plot, normalized such that t=0 represents the cell’s first contact with the interface. Bars are color coded to match (A). **(C-D)** The final outcomes at the end of 45 minutes of imaging *Naegleria* (C) or *Dictyostelium* (D) were quantified as a percentage. Each dot represents a single trial, while bars show the average.

To determine if the tendency of *Naegleria* to enter channels is unusual, we again compared *Naegleria* to *Dictyostelium* (**Fig. 3D, S6B**). Unlike *Naegleria*, *Dictyostelium* cells were fairly inefficient at entering channels, with at most 51.3 ± 33.4% entering 6 μm channels (**Fig. 3D**). *Dictyostelium* also showed high rates of disengagement (>25%), independent of channel width. These data highlight a stark difference between *Naegleria* and *Dictyostelium*: while *Naegleria* enters channels at high rates and readily disengages from too-narrow openings, *Dictyostelium* is less efficient at entry, and has higher disengagement for all channel widths.

### *Naegleria* cells readily explore complex, three-dimensional environments

*Naegleria’s* tendency to crawl into channels is consistent with its presence within layers of silt and sediments at the bottom of lakes and ponds (Stahl and Olson 2020). To assess whether their behavior under confinement reflects how they move in natural environments we presented *Naegleria* amoebae with granular media that better mimics lake sediments. To directly visualize *Naegleria’s* motility in these media, we took advantage of the transparency of Carbopol hydrogel microparticles. These particles are composed of randomly crosslinked polymer chains that are strongly hydrophilic and swell in liquid, resulting in a matrix of densely-packed, optically transparent liquid-infused particles around ∼5 μm in diameter, a similar size to silt (Friedman and Sanders 1980). To characterize the mechanical properties of the Carbopol matrix, we conducted rheological measurements (**Fig. S7**) that showed that Carbopol exhibits yield stress behavior, behaving as an elastic solid under small stresses and rearranging and flowing like a viscous fluid when imposed stresses exceed the yield stress (*σ*y). Importantly, we can tune yield stress by adjusting the Carbopol particle concentration (**Fig. S4**), allowing us to mimic the variability of natural silts, which display a wide range of yield stresses (Berlamont et al. 1993).

To explore if and how *Naegleria* moves within these environments, we mixed cells with Carbopol and imaged them at high magnification (**Fig. 4A**). We observed cells within the matrix crawling using a combination of lamellar protrusions and blebs, with cells frequently switching between these two modes or using both simultaneously (**Fig. 4A, Video S3**). To determine how the mechanical properties of the matrix could influence cell crawling, we introduced cells into Carbopol matrices of three different concentrations and imaged the cells as they moved through the interstitial pore space between particles in three dimensions. The mass fraction of dry hydrogel particles used to prepare each matrix was 0.3%, 0.6%, or 1.0%, resulting in jammed packings of the swollen particles that are increasingly dense, with progressively smaller interstitial pores (Bhattacharjee and Datta 2019b, 2019a) and higher yield stresses (34.1 Pa, 103.5 Pa, and 130.3 Pa, respectively). *Naegleria* amoebae readily migrated without any apparent directional bias through 0.3% and 0.6% Carbopol matrices, but had limited motion and became trapped in 1.0% Carbopol (**Fig. 4B**), indicating that *Naegleria’s* motility is strongly influenced by the mechanical properties of the matrix. Three-dimensional tracking of individual cells also showed that the maximum cell speed decreased with increasing Carbopol density (19.5 μm/min in 0.3%, 12.6 μm/min in 0.6%, and 3.5 μm/min in 1.0% Carbopol), (**Fig. 4B-C**), mirroring the slowing we observed in narrower channels. We also observed that cells in higher density Carbopol matrices become *less* persistent, perhaps due to becoming stuck in smaller pore spaces, trapped by an abundance of possible paths (Ron et al. 2024; Renkawitz et al. 2019), and/or due to stiffer physical environments (Truszkowski et al. 2023).

**Figure 4.**
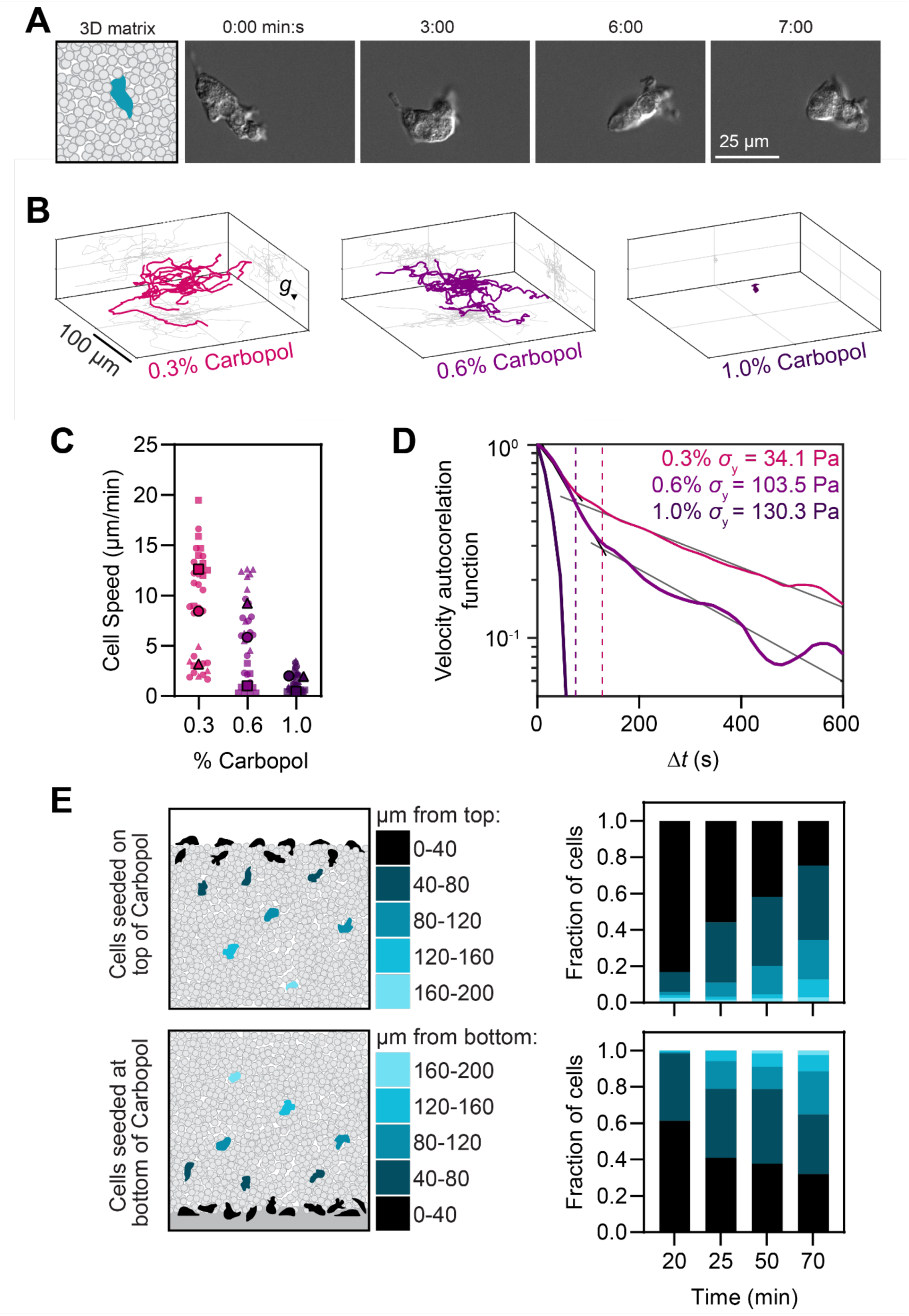
*Naegleria* readily enters and crawls through Carbopol particle matrices. (A) *Naegleria* cells were suspended in a 0.45% carbopol matrix, and imaged in DIC at high magnification. A cartoon showing the experimental setup is on the left, and select DIC images on the right. **(B-D)** Cells were imaged as in A, but with 0.3, 0.6, or 1.0% carbopol at low magnification. Cell positions were tracked in x, y, and z over time. Representative tracks from one of three replicates are shown (**B**), while cell speeds are displayed as a SuperPlot (**C**) with small symbols representing speeds of individual cells, and large symbols representing experiment-level averages, with shape coordinated by trial. The Velocity Autocorrelation was determined of each trajectory and the average is shown in (**D**), which is plotted with grayscale lines indicating the two slope regimes for the 0.3 and 0.6% carbopol, and dashed lines indicating the time interval at which this regime changes. **(E)** Cells were seeded on top of a layer of carbopol (top left), or overlaid with Carbopol (bottom left), and 200 μm z stacks were acquired over a 70 minute period. The number of cells in each frame was normalized to the total number of cells in the sample at that time, and plotted as a histogram based on z position (right). Histograms show one representative experiment.

Because the traditional persistence analysis used for our prior experiments cannot reliably quantify persistence in three dimensions, we calculated the velocity autocorrelation function (VACF) of cells migrating through Carbopol, where faster VACF decay (steeper slopes) indicates less persistent movement. We found that VACF decreases with increasing Carbopol density (**Fig. 4D**), indicating that *Naegleria* are more persistent in lower yield stress matrices. This result contrasts with the higher persistence of *Naegleria* confined in channels, indicating a fundamental difference in cell behavior in these different environments. This analysis also revealed distinct phases in both 0.3 and 0.6% Carbopol, with steeper slopes at short (0-75 seconds for 0.6% Carbopol and 0-127 seconds for 0.3% Carbopol) compared to long timescales (>75 seconds for 0.6% and >127 seconds for 0.3% Carbopol,**Fig. 4D**). These results are consistent with our observations of non-migratory cells probing their surroundings as well as cells undergoing productive movement over longer periods similar to white blood cells migrating through collagen (Fritz-Laylin et al. 2017).

Having established that *Naegleria* amoebae crawl within Carbopol matrices, we next wondered whether they would voluntarily enter these complex, three-dimensional environments. We therefore seeded *Naegleria* cells on top of Carbopol matrices and tracked cell concentration at different depths over time. Consistent with *Naegleria’s* affinity for crawling into channels, >75% of cells disperse at least 40 μm into the granular matrix, with >30% reaching depths of 80 μm or greater over the course of the experiment (**Figure 4E**). Because this approach can only reveal downward burrowing behavior, we also overlaid gel on top of amoebae to assess their ability to climb upwards into the matrix. Again, we observed amoebae readily entering the gel, with >60% migrating at least 40 μm vertically from the coverslip, and >30% migrating 80 μm or more. Taken together, these data indicate that *Naegleria* readily enters and navigates complex, three-dimensional environments using multiple modes of cell migration.

## DISCUSSION

Here we show that *Naegleria gruberi* amoebae modulate their behavior in an environment-dependent manner. While crawling across flat surfaces, *Naegleria* uses a dynamic combination of actin-rich lamellar protrusions and blebs. When confined in narrow channels, however, *Naegleria* amoebae switch to exclusively bleb-driven motility. Unlike *Dictyostelium*, which frequently disengages from channel interfaces, *Naegleria* probe channel openings until they enter. Once inside, amoebae maintain rapid, unidirectional movement for over a millimeter. We also show that *Naegleria* readily burrows into sediment-like matrices using both blebbing and lamellar protrusions. Collectively, these findings reveal that *Naegleria* amoebae explore and navigate diverse microenvironments using multiple, distinct migration strategies.

We also show that the geometry of *Naegleria*’s microenvironment shapes how *Naegleria* crawls: Cells confined between two flat coverslips use a mixture of blebs and lamellar protrusions, while cells crawling through rectangular channels only use blebbing. One possible explanation for this difference is that cells sandwiched between coverslips are confined on two sides, while cells in narrow channels experience higher degrees of confinement, perhaps enough to trigger the switch to exclusively bleb-based motility. The geometry of the channels may also enable additional mechanisms of migration, such as versions of a nuclear piston (Petrie et al. 2014) and/or an osmotic engine (Stroka et al. 2014). *Naegleria* cells also crawled faster and more persistently in channels than on flat surfaces, even faster and and more persistent than *Dictyostelium* in the same environments. While it is tempting to speculate that blebbing is directly responsible for *Naegleria*’s rapid movement through channels, studies of other species suggest a more complex, indirect relationship where blebbing motility can be faster, slower, or the same speed as lamellar-based motility depending on the degree of confinement, surface coatings, and media (Ruprecht et al. 2015; Liu et al. 2015; Yoshida and Soldati 2006; Srivastava et al. 2020; Te Boekhorst et al. 2022). Although cell speed and persistence are clearly modulated by many environmental factors (Paulson et al. 2025; Stinson et al. 2025; Petrie et al. 2012; Gaertner et al. 2022; Bodor et al. 2020), the increase in *Naegleria*’s speed and persistence in channels may be explained, at least in part, by directional polarity imposed by channel walls restricting lateral protrusions and favoring forward-directed ones.

We also explored how *Naegleria* amoebae migrate through silt-simulating matrices with complex, three-dimensional geometries and found that these cells crawl using both blebs and lamellar protrusions. This behavior is more similar to cells migrating across flat surfaces or sandwiched between two coverslips than cells migrating through rectangular channels. Although unexpected, given that both channels and matrices force cells to squeeze around obstacles to move, this result can be explained by differences in the geometry of these two microenvironments. Unlike the uniform compression experienced by cells in channels, cells in matrices are non-uniformly compressed within the variable pore spaces between Carbopol particles. Such variable compression may be insufficient to promote constant blebbling-only motility. Additionally, cells in higher density Carbopol matrices become *less* persistent, perhaps due to becoming stuck in smaller pore spaces, or trapped by an abundance of possible paths (Ron et al. 2024; Renkawitz et al. 2019). Finally, similar to previous work in *Naegleria* and other organisms (Uwamichi et al. 2023; Selmeczi et al. 2008), we found that longer time scales correlate with increased persistence in complex environments, suggesting *Naegleria* cells may retain memory of past orientation.

Finally, we show that *Naegleria* amoebae readily enter 4 to 8 μm channels in the absence of chemoattractant and crawl into Carbopol matrices from both above and below. Moreover, *Naegleria* cells also inserted protrusions at high frequency into channels too narrow to enter, suggestive of a frustrated ability to enter confined spaces. This tendency to enter confinement without inducement—or “claustrophilia”—is more pronounced in *Naegleria* than in *Dictyostelium*, a difference that could reflect adaptation to their natural habitats. While both species hunt bacteria in complex three-dimensional environments, *Dictyostelium* inhabits forest soil (Raper and Rahn 1984), while *Naegleria* is typically isolated from silt and leaf litter at the bottom of ponds (Fulton 1970). Differences in the spatial organization and physical properties of these microhabitats could make claustrophilia more beneficial to *Naegleria*.

Taken together, these data show that *Naegleria* exhibits environment-dependent migration strategies that, while normally benign, in the proper context could promote pathogenesis., In three-dimensional matrices resembling silt, cells undergo random walks with directional memory at longer timescales—a pattern consistent with exploration and foraging. These amoebae also actively seek confinement, exhibiting pronounced claustrophilia, and crawl rapidly with near-perfect persistence across millimeter-scale distances when confined to tight channels. We propose that these three behaviors—memory-guided exploration, claustrophilia, and persistent one-dimensional crawling—are adaptations for life in pond sediments that enable *Naegleria* to detect, probe, and navigate tight microenvironments between sediment particles that harbor bacterial prey. During human infection, however, these same adaptations may inadvertently promote *N. fowleri* pathogenesis: directional memory would allow exploration of the sinus cavity, claustrophilia could draw amoebae into narrow spaces between olfactory axons, and persistent crawling in narrow channels may facilitate rapid transit to the brain. In this way, behaviors optimized for survival in pond sediments become mechanisms of deadly invasion in human hosts.

## METHODS

### Cell culture

#### Naegleria gruberi

Amoebae (strain NEG-M, a gift from Chandler Fulton) were cultured as previously described (e.g. (Velle et al. 2023). Briefly, cells were thawed from liquid nitrogen or -80 freezer stocks, and grown in M7 media (10% FBS + 45 mg/L L-methionine + 5 g/L yeast extract + 5.4 g/L glucose + 2% (v/v) M7 Buffer (18.1 g/L KH2PO4 + 25 g/L Na2HPO4)) in 25 cm^2^ plug-seal tissue culture treated flasks at 28 °C. Cells were split into fresh flasks once they reached ∼70% confluency by 1:20-1:40 dilutions approximately 3 times per week. Cells were used for experiments between passages 3 and 20.

#### Dictyostelium discoideum

Cells (strain Ax2-ME, cloned and expanded from AX2-RRK DBS0235521 obtained from the Dictybase stock center (Fey et al. 2013)) were grown axenically in HL5 Media (Formedium, Norfolk, UK) supplemented with streptomycin (300 μg/mL) and ampicillin (100 μg/mL) on tissue culture-treated plastic 100 mm or 150 mm plates at 21 °C and maintained at 70% confluency by sub-culturing.

### OneStep fixing and staining

The experiments shown in **Fig. 1A**, **Fig. S2**, and **Fig. S4A** were completed following our published protocol (Velle et al. 2021), with the following specifications: for **Fig 1A**, 100-150 μL of NEG-M cells from a healthy flask (50-70% confluent) were seeded into a 96 well glass-bottom plate (Brooks) to achieve a confluency of ∼30%, and allowed to settle for a few minutes. The media was then removed by gently pipetting, and 150 μL 2 mM Tris, pH 7.2 were added. Cells were then imaged live using DIC and a 100x oil objective, at a rate of 2 sec/frame. After a few minutes of imaging, 150 μL of 2x OneStep solution were added (130 mM sucrose, 50 mM sodium phosphate buffer (pH 7.2), 3.6% PFA, 0.005% NP-40 alternative, 66 nM AlexaFluor-488 labeled phalloidin, 2 μg/ml DAPI), and cells were imaged using DIC, GFP, and DAPI settings until staining plateaued (see additional details in “Microscopy”). For **Fig. S2** and **S4A**, ∼5,000-7,500 cells were incubated in each well of a dish with channels (see below for details) before adding OneStep solution. Fig. S4A used 10 μM Hoechst 33342 instead of DAPI.

### Cell confinement between coverslips

Cells were confined using previously developed methods (Le Berre et al. 2014) with a dynamic cell confiner system commercially available from 4Dcell that included a PDMS suction cup (reusable), a confinement coverslip with 5 μm PDMS spacer pillars (single use), and a vacuum pump. The PDMS suction cup and confiner coverslip were each incubated in 2 mM Tris for at least 30 minutes prior to the experiment to allow equilibration. *Naegleria* cells were seeded in a 6-well glass-bottom plate (Cellvis) to <20% confluency and allowed to adhere, then media was replaced with 2 mM Tris. Cells were imaged by phase contrast for 20 minutes at a rate of 5 sec/frame. Cells were confined by lowering the confinement coverslip by gradually decreasing the vacuum pressure from -30 mbar to -100 mbar over about 30 seconds. Cells were then imaged for the following 20 minutes (see “Data analysis and statistics” for additional information on cell quantifications).

### Cell confinement in channels

Microchannel dishes (MC004) were purchased from 4Dcell: each dish contains 9 wells with 6 microchannel runs that connect adjacent wells. A fresh dish was used for each experimental replicate, and was prepared by rinsing thrice with ∼3 mL MilliQ water and pipetting up and down in each well on the last wash. Dishes were then rinsed thrice in ∼3 mL of 2 mM Tris (pH 7.2), and incubated for at least 30 minutes to allow the PDMS to equilibrate. After this incubation, the Tris was aspirated from the dish and wells, and 3 mL of fresh Tris were added to fully submerge the PDMS. 2 mL of Tris were then removed so the liquid level dropped below the well level. *N. gruberi* cells were prepared by taking a sample of cells from an actively growing flask, spinning them down at 1500 RCF for 90 s, and resuspending in Tris to a final concentration of 5E5 cells/mL immediately before adding them to a well. Fresh cells were prepared for each width tested. For each width, 10 μL of Tris were removed from a well, and 10 μL of amoebae were added for a total of 5,000 cells per well. A coverslip was placed on top of the well to prevent the meniscus from interfering with imaging. Cells were imaged using a 20x DIC objective for 45 minutes with 2 seconds between frames (see additional imaging details in “microscopy”).

### Cell confinement in carbopol matrices

To create porous 3-D matrices, dry Carbopol granules of randomly crosslinked acrylic acid/alkyl acrylate copolymers (Carbomer 980, Ashland) were dispersed directly into 2 mM Tris to make 0.3, 0.45, 0.6, or 1% (weight/volume) Carbopol suspensions. The suspensions were mixed for at least 12 hours to ensure a homogeneous distribution and the pH subsequently adjusted to 7.4 by the addition of 10M NaOH. Carbopol was briefly spun down at 4000 rpm to remove air bubbles before each use. Yield-stress quantification was performed using an Anton Paar MCR 502 rheometer using forward and reverse strain rate sweeps.

To track *Naegleria* motility in carbopol matrices (**Fig. 4A-B**), 0.5-1 mL of cells from a growing flask (∼70% confluent) were spun down at 1500 RCF and pellets were resuspended in 1 mL of Carbopol using a wide gauge needle and syringe. This mixture of Carbopol and cells was then transferred to a glass-bottom petri dish (with a packing thickness of approximately 1 mm), and for cell tracking experiments (**Fig. 4B-D**), a volume of 850×850×110 µm^3^ was imaged for 1 hour, every 14.7 seconds.

To test the ability of cells to enter carbopol matrices (**Fig. 4E**), cells were either seeded on the bottom of a glass imaging dish with carbopol then added on top, or the carbopol was added to the dish, and then cells were added to the top, and a 200 µm thick *z* stack was captured at 20, 25, 50, and 70 minute timepoints. (See “Microscopy” for imaging details and “Data analysis and statistics” for additional information on cell quantifications).

To quantify the mechanical response of the hydrogel matrices, we measured their flow curves using an Anton Paar MCR 502 rheometer. Approximately 2–3 mL of each sample was loaded into a 1 mm gap between 50 mm-diameter parallel plates. To prevent wall slip, we used a roughened upper plate and affixed sandpaper to the lower plate with double-sided tape. At vanishing shear, the matrices behave as jammed solids. To probe their fluidization under shear, we vary the shear rates from 0.001–200 s^-1^ and measure the resulting shear stress in the matrix (**Fig. S7**). At low shear rates, the shear stress remains nearly constant and independent of shear rate, consistent with a finite yield stress. Beyond this regime, the samples fluidize and the stress exhibits a power-law dependence on shear rate. The full flow curves are captured well by the Herschel–Bulkley model (solid lines in **Fig. S7**). We find that as the volume fraction of the carbopol increases, the matrix becomes more solid and the yield stress increases. Using this approach, we produce transparent viscoplastic hydrogels with tunable yield stresses between 34.1 and 130.3 Pa.

### Naegleria and Dictyostelium size analyses

To determine the footprint of whole cells and nuclei in Fig. S4B-C, approximately 5x10^3^ *D. discoideum* cells or 1.7x10^3^ *N. gruberi* cells in 1 mL of M7 media were fixed in suspension by adding an equal volume of fixative (3.6% PFA, 130 mM sucrose, 50 mM sodium phosphate buffer (pH 7.2)) for 10 minutes. Cells were then transferred to wells of a PEI-coated 96-well glass-bottom plate, and allowed to adhere for 10 minutes. Cells were rinsed once with PEM (100 mM PIPES + 1 mM EGTA + 0.1 mM MgSO4), permeabilized in 0.1% NP-40 alternative supplemented with 6.6 nM AlexaFluor-488 phalloidin, rinsed with PEM, then labeled for 35 minutes with 66 nM AlexaFluor-488 phalloidin and 1 μg/ml DAPI. Cells were rinsed once in PEM, then imaged. 30 cells of each species were randomly selected for analysis in Fiji, excluding cells that were multinucleated or in contact with other cells. The “freehand selection” tool was used to outline the whole cell based on phalloidin staining and also to outline nuclei based on DAPI. Areas were quantified using the “measure” function.

### Microscopy

The microscopy data shown in **Fig. 1A, 2D, 3A, 4A, S1A, S2, S4B,** and **Video S1,** were collected using a Nikon Ti2E, equipped with a Plan Apochromat λ D 100x 1.45 NA oil objective (**Fig. 1A, S1A,** and **Video S1**) or a Plan Apochromat λ 20x 0.75 NA air objective (**Fig. 2D, 3A, S2,** and **S4B**), a Photometrics Prime BSI Express camera, and a SOLA light engine Gen III (for fluorescence) or Diascopic illumination (for DIC). The microscope was controlled through NIS-Elements AR software. DAPI was imaged using a 378/52 nm excitation filter and 447/60 nm emission filter, and AlexaFluor-488 Phalloidin was imaged using 466/40 nm excitation and 525/50 nm emission filters.

The phase contrast microscopy shown in **Fig. 2A** was performed using a Nikon Ti2E, equipped with a Photometrics Prime BSI Express camera and a Plan Fluor 10x Ph1 NA 0.3 objective, controlled using NIS elements software.

The microscopy in **Fig. S2** was performed using a Nikon Ti2 microscope equipped with a a Spectra III/Celesta light source (at 50–100% power with excitation wavelengths of 405, 477, 546, and 638 nm), a Crest spinning disk (50 μm), a Plan Apo λ 100x oil objective (1.45 NA), and a Prime 95B CMOS camera, controlled through NIS Elements software.

*Dictyostelum* cells from experiments shown in **Fig. 2** were imaged using a ZEISS Observer 7 running Zen Blue (ZEISS) software, equipped with an Orca Flash CMOS camera (Hammamatsu, Japan) using a 20X Plan Achromat 0.75 N.A. objective for low magnification, or a Plan Achromat 63x/1.40 N.A. objective for high magnification.

To track *Naegleria* motility in 3D granular media (**Fig. 4B-E**), 1 mL of jammed hydrogel media containing cells were confined at the bottom of a sealed glass-bottom petri dish resulting in a packing thickness of approximately 1 mm. Cells were imaged using a Nikon A1R inverted laser-scanning confocal microscope equipped with a temperature-controlled stage set at 28 °C, and a 10x, 0.45 NA objective. For data in **Fig. 4B-D**, a volume of 850 x 850 x 110 µm^3^ was scanned every 14.7 s, capturing bright-field images with a *z* step of 10 µm. For the experiments presented in **Fig. 4E**, large images were stitched from the captured bright-field frames in *x*-*y* plane yielding a field of view of 2400 x 2400 x 200 µm^3^ at different heights with a z step of 20 µm. The volume of interest was scanned every 65 s.

### Data analysis and statistics

#### Scoring blebs and lamellar protrusions

The quantification shown in **Fig. 1C** was completed using the live imaging data from experiments shown in **Fig. 2A** and **Fig. 2F/3A**. Two movie clips, each consisting of two consecutive frames, were selected at random from each movie, one clip pre confinement and the second post confinement. Every cell that had at least one obvious lamellar protrusion or obvious bleb was scored—cells without clear protrusions were excluded from the analysis. For confinement between coverslips, 35-128 cells were scored for each of four trials), and for confinement in channels, 8-66 cells were scored for each of three trials.

#### Tracking speed and persistence

For confinement coverslip experiments (**Fig. 2** **A-C**), the approximate center of 30 randomly selected cells that were in the field of view for the entire imaging period were tracked before and after confinement in Fiji (Schindelin et al. 2012) using the mTrackJ plugin (Meijering et al. 2012). Speed was calculated by dividing the path length by the imaging time (20 minutes). Directional persistence was calculated by dividing the maximum displacement from the starting point by the total path length. The averages from four independent biological replicates were used for paired t tests comparing pre and post confinement conditions. Graphs were generated using GraphPad Prism in the SuperPlot style (Lord et al. 2020).

Measurements of speed and persistence in channels for *Naegleria* and *Dictyostelium* (**Fig. 2** **F-I**) were also completed using Fiji and the mTrackJ plugin. For these experiments, cell tracking began once a cell had fully entered a channel, and ended either when the cell left the field of view, or when the movie ended. Because the channel entry rate limited the sample size, up to 30 *Naegleria* cells were tracked for 3 trials and up to 12 *Dictyostelium* cells were tracked for 4 trials. Cells were tracked from the rear edge to account for fluctuations that would sometimes occur at the leading edge that affected the position of the “center” of the cell. Speed and persistence were then calculated from the tracking data as described above. Ordinary one-way ANOVAs and Tukey’s multiple comparisons tests were used to compare the experiment level averages across channel sizes and to unconfined cells from the same experiment.

#### Analyzing interactions with microchannels

For the analysis shown in **Fig. 3** **B-D** and **Fig. S6**, up to the first 21 *Naegleria* or the first 13 *Dictyostelium* cells to make contact with the microchannel interface were tracked and categorized over time. Cells were categorized as either “contacting” the interface once it was clear some part of the cell was touching the PDMS surface, “protruding” once any part of the cell crossed into one or more channels, or “inside” once the cell was entirely confined in a channel. Each cell was tracked for categories either until it left the field of view, or the movie ended. These categorical tracks were then normalized such that the initial contact time was set to 0, and data were graphed as horizontal stacked bar plots in GraphPad Prism. The category of each cell at the end of the video (or at the end of its track) was then used to plot the outcomes as a percentage for each of the 3 experimental replicates for each species (**Fig. 3C-D**).

#### Quantifying cell motility in carbopol matrices

To track the cells in 3D, the images were preprocessed to remove noise, after which the ImageJ Plugin TrackMate (Tinevez et al. 2017) was used to reconstruct the 3D matrix of cells by identifying the objects and extracting the coordinates in 3D and then finding the tracks. MATLAB was used to perform statistical analysis on the obtained trajectories.

To measure the velocity autocorrelation function, we used

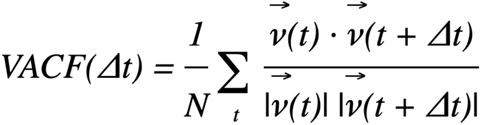

where v(t) is the velocity of the amoeba at time t, |v(t)| is the speed, N is the number of time steps, and \delta t is the lag time.

## ACKNOWLEDGEMENTS

This work was supported by The National Institute of General Medical Sciences of the National Institutes of Health under Award Numbers: R00GM147656 to K.B.V., 1R15GM143733-01 to M.E., and GM143039 to L.K.F.-L.; and The National Science Foundation under Award Numbers: DMR-2011750 to M.R., and CBET-1941716 and DMR-2011750 to S.S.D. This work was also supported by a Research Award from Amazing Aven’s Quest for Amoeba Awareness to K.B.V., the Amherst College Promise Campaign (awarded to W.S.C.), the Amherst College Greg Call Research Support Program (awarded to H.S.K.), the Camille Dreyfus Teacher-Scholar Program (awarded to S.S.D) and the Pew Biomedical Scholars Program (awarded to S.S.D). L.K.F.-L. is an Investigator of the Howard Hughes Medical Institute and a Fellow of the Canadian Institute for Advanced Research Fungal Kingdom: Threats and Opportunities Program. The authors wish to thank Iain Patten for valuable feedback and suggestions on the manuscript.

## SUPPLEMENTAL VIDEO LEGENDS

**Video S1. Unconfined *Naegleria* form a range of blebs and lamellar protrusions. (A)** The top cell shown in Fig. 1A forms lamellar protrusions to crawl. In situ fixing and staining shows these protrusions contain actin polymer. The scale bar indicates 10 μm. **(B)** The bottom cell from Fig. 1B forms blebs, which lack actin polymer. Flashing lines indicate actin arcs. The scale bar is 10 μm. **(C)** A cell forming a dynamic mixture of blebs and lamellar protrusions was imaged live, then fixed and stained in situ. Actin polymer staining shows an actin-rich lamellar protrusion towards the bottom of the cell, and an actin arc and cortex consistent with blebbing towards the top of the cell. **(D)** two examples of cells that switch between blebs (labelled with a B) and lamellar protrusions (labeled with an L).

**Video S2. *Naegleria* amoebae favor interactions with surfaces and channels.** Videos show cells interacting with 8, 6, 4, and 2 μm channels, and with a PDMS surface lacking channels. Data are from experiments shown in Fig. 3.

**Video S3. *Naegleria* use a mixture of blebs and lamellar protrusions to crawl in Carbopol** Two representative cells crawling through 0.45% Carbopol are shown, with a few examples of blebs (labelled B) and lamellar protrusions (labelled L). The first cell in the video is the same cell shown in **Fig. 4A**.

## SUPPLEMENTAL FIGURES

**Figure S1.**
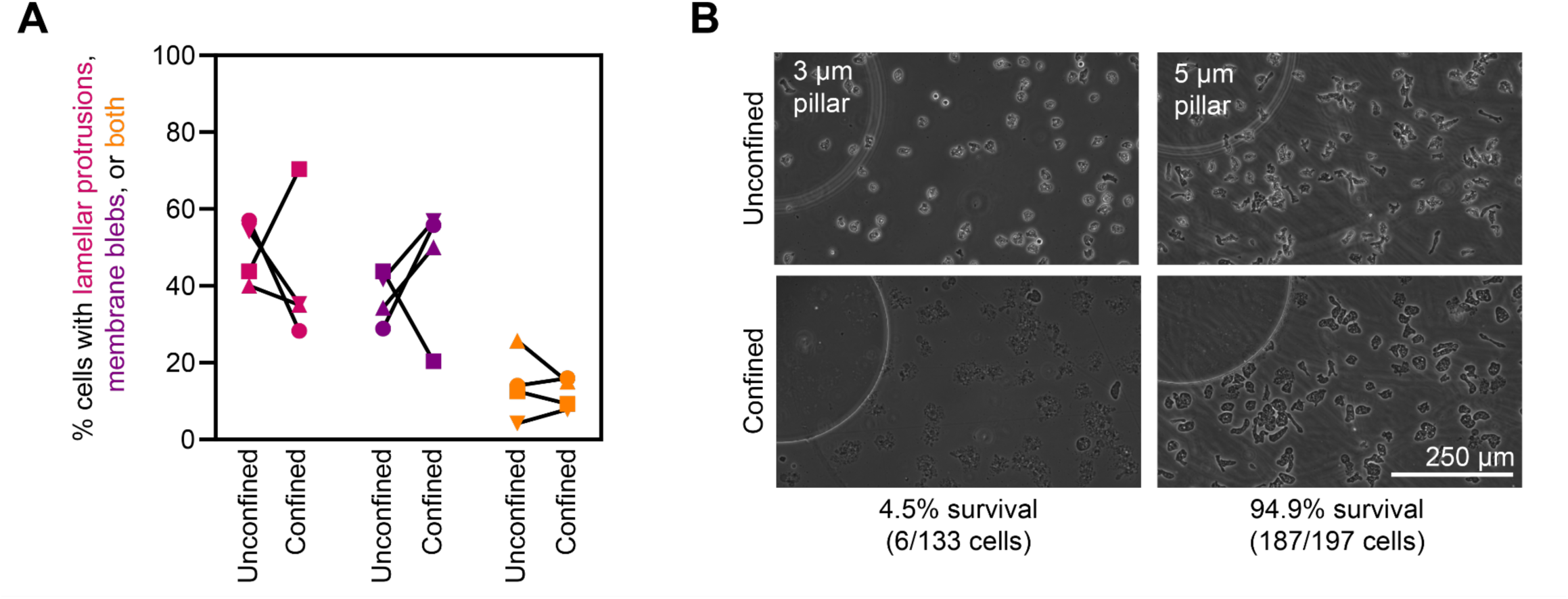
Confinement between coverslips leads to variable phenotypes. **(A)** The data from individual trials using confinement coverslips which were combined in Fig. 1C are shown by replicate (coordinated by shape). **(B)** 3 μm and 5 μm pillar sizes were tested to determine which size led to optimal confinement between coverslips. Confinement to 3 μm caused the vast majority of cells to burst. Images show the same representative area before and after confinement.

**Figure S2.**
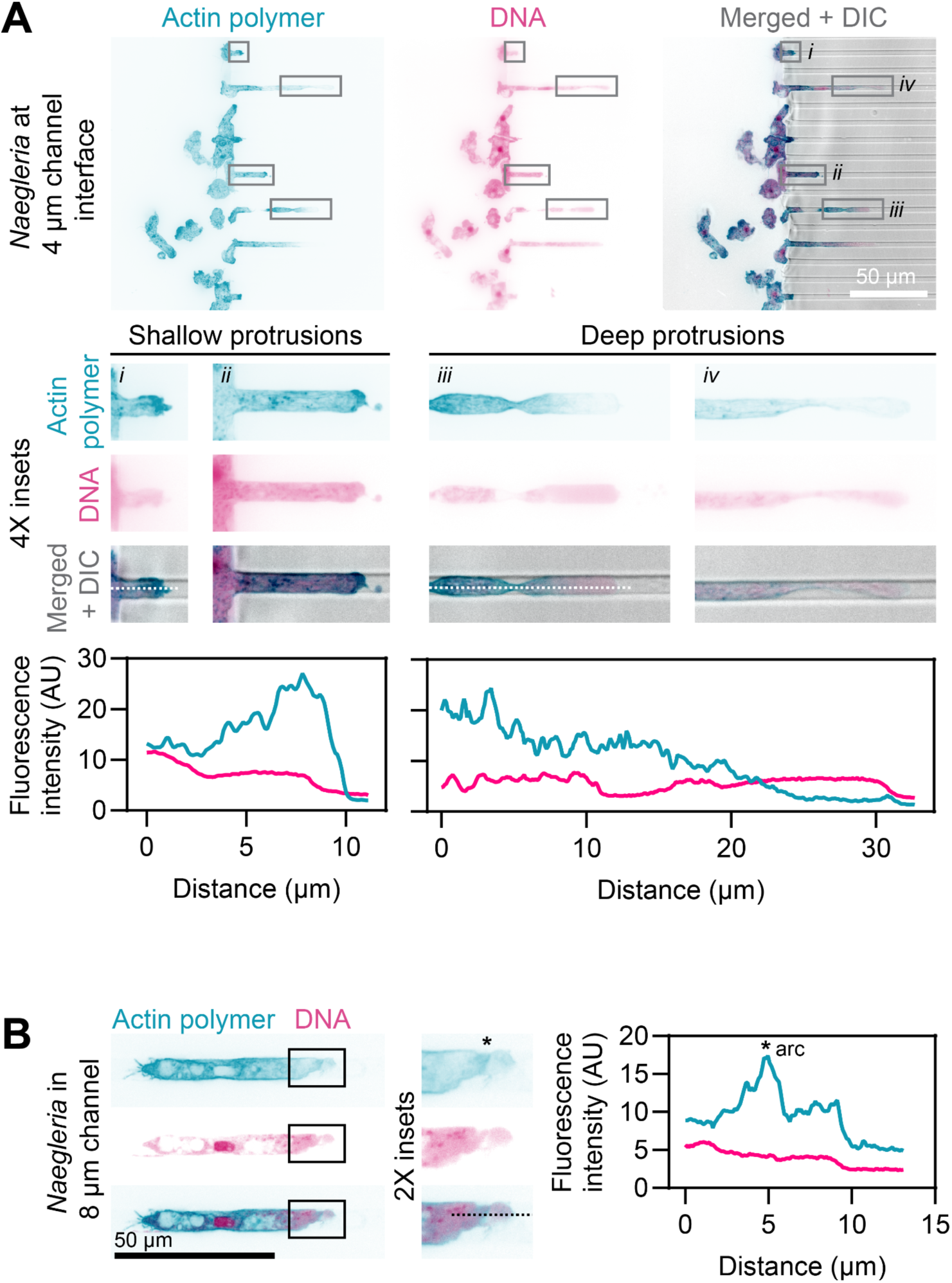
*Naegleria* cells switch from actin-rich protrusions to blebs when entering channels. **(A)** *N. gruberi* cells at the entrance of 4 μm channels were fixed and stained *in situ* for actin polymer (teal) using phalloidin and for DNA (magenta) using DAPI. Insets (middle row) show examples of shallow or deep protrusions. Linescans (bottom row) show actin polymer (teal) or DNA (magenta) staining intensity along the white dotted lines indicated in panels i and iii. **(B)** Cells were fixed and stained as in (A), but in an 8 μm channel. The leading edge of the cell is shown as an inset, with an actin arc indicated with an asterisk. A linescan along the black dotted line shows fluorescence intensity of actin polymer (teal) and DNA (magenta), with the asterisk corresponding to the position of the actin arc.

**Figure S3.**
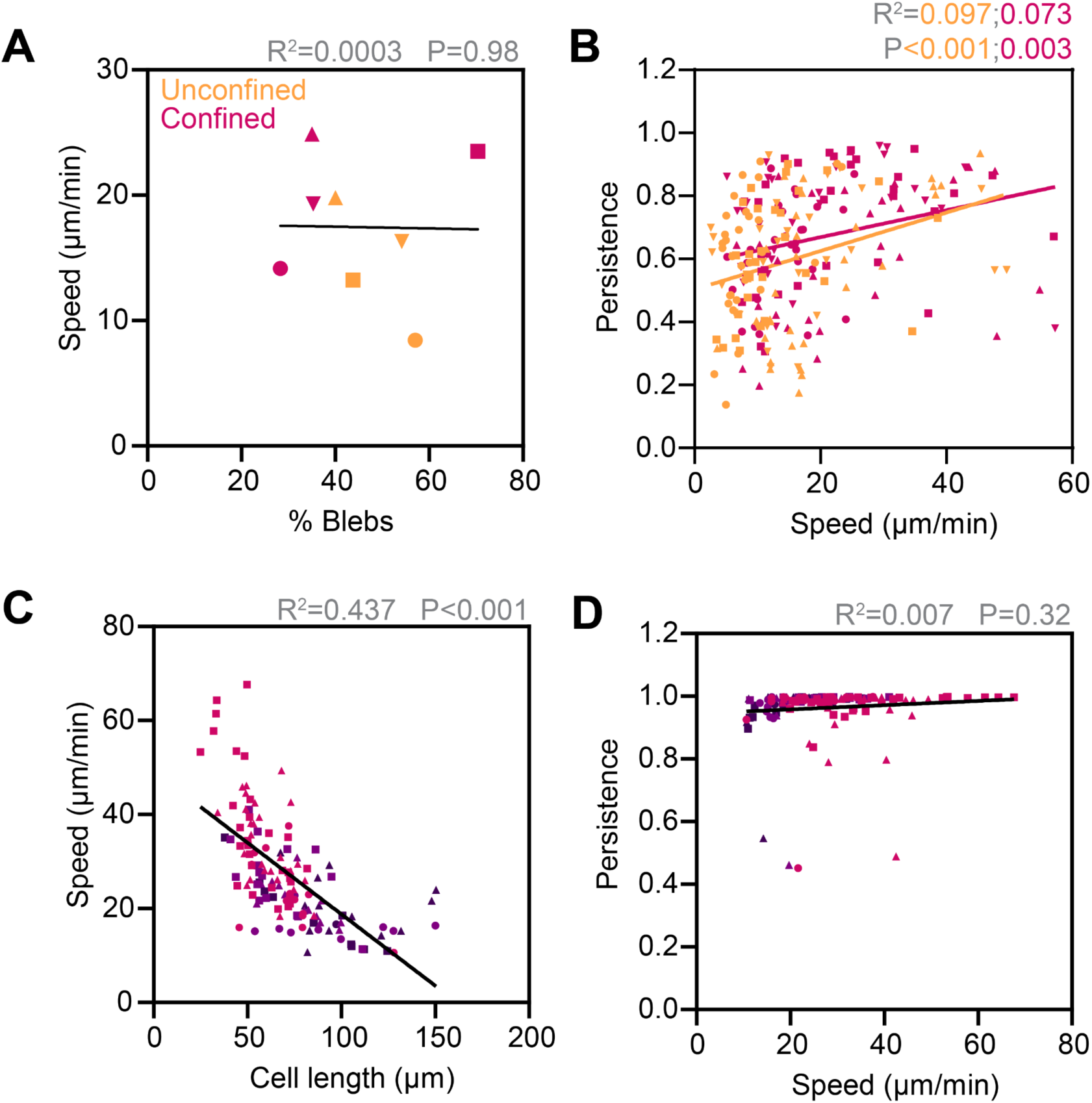
Cell speed is negatively correlated with cell length. **(A)** Data from experiments shown in Fig. 1C and Fig. 2B were plotted to compare average speeds and the percent of the cell population that used blebs. Data from unconfined populations (orange) and cells confined between coverslips (pink) were pooled for analysis. **(B)** Data from Fig. 2B-C were plotted to compare speed and persistence for each cell. Unconfined (orange) and confined (pink) populations were each analyzed separately. Symbol shapes are coordinated by trial and match A. **(C)** The lengths of cells from Fig. 2F in 8 μm channels (pink), 6 μm channels (light purple), or 4 μm channels were plotted against their speeds. **(D)** Data from Fig. 2F and 2H were plotted to compare speed and persistence for each cell. Data in all panels were analyzed by simple linear regression.

**Figure S4.**
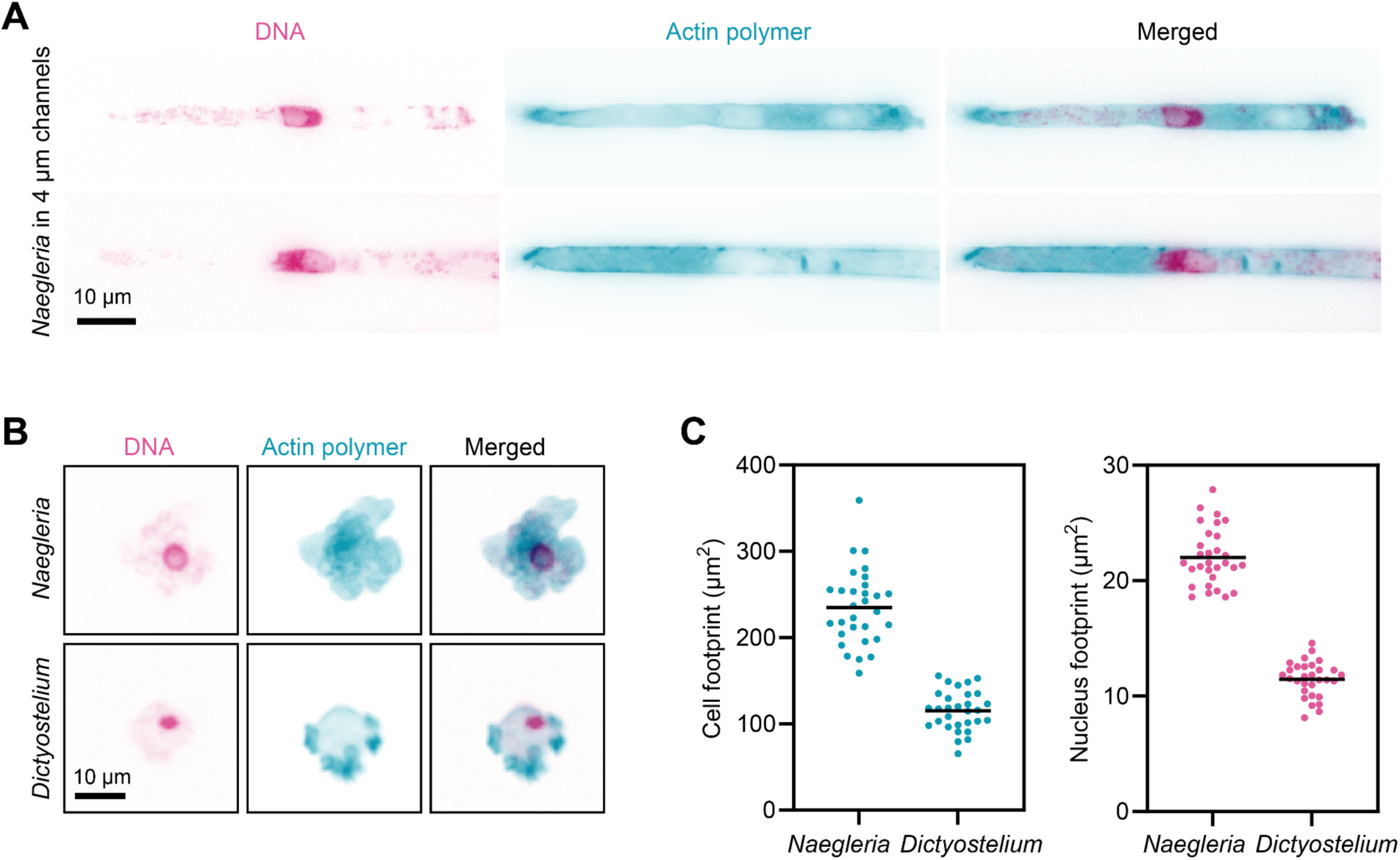
*Naegleria*’s nuclei deform in channels. **(A)** *N. gruberi* cells crawling in 4 μm channels were fixed and stained with Hoescht to visualize DNA (magenta), and phalloidin to label actin polymer (teal). Two representative cells with clear nuclear squishing are shown. **(B)** *N. gruberi* and *D. discoideum* cells were fixed then stained with DAPI to visualize DNA (magenta) and phalloidin to label actin polymer (teal). A representative cell for each species is shown. **(C)** Actin staining was used to outline the cell footprint to quantify cell areas, and DNA staining was used to quantify nuclear footprints. Each symbol represents an individual cell. The mean is indicated with a black line. N=30 cells.

**Figure S5.**
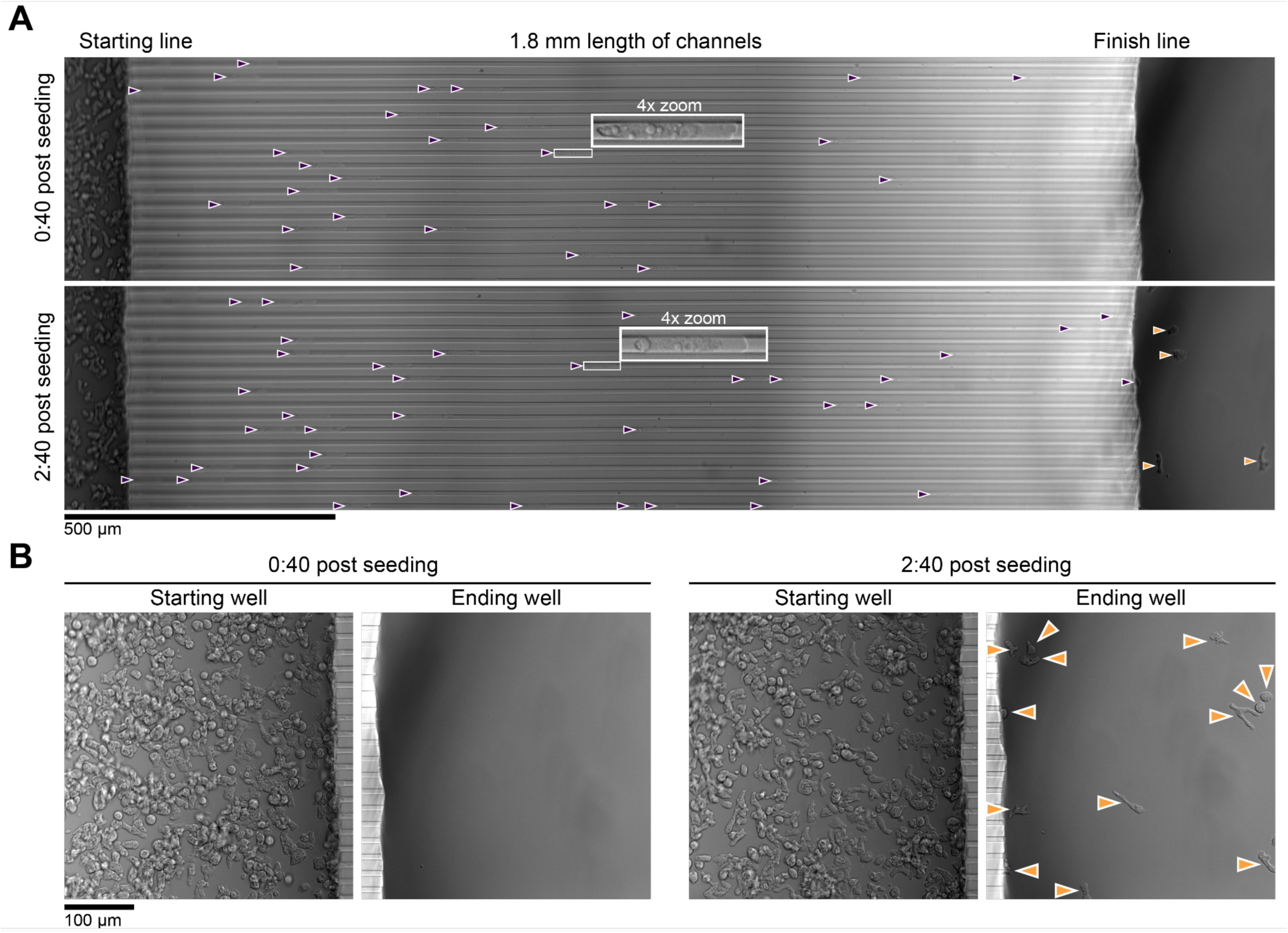
*Naegleria* amoebae can successfully complete a 1.8 mm dash. **(A)** *Naegleria* amoebae were seeded into a single well (left) of a dish with 8 μm channels, and imaged for 2 hours, starting 40 minutes post seeding. Images cover the full 1.8 mm run and the adjacent starting and ending wells. A single focal plane where the channels are in focus is shown for each timepoint. 4X insets show magnified amoebae from an optimized focal plane. Cells in channels are noted with purple arrowheads, cells that traversed the entire run length to reach the next well are shown with orange arrowheads. **(B)** Higher magnification views at optimized focal planes from the timepoints in A are shown for the starting well (where cells were seeded) and the ending well (where cells crawled). Cells that made it past the finish line are labeled with an orange arrowhead.

**Figure S6.**
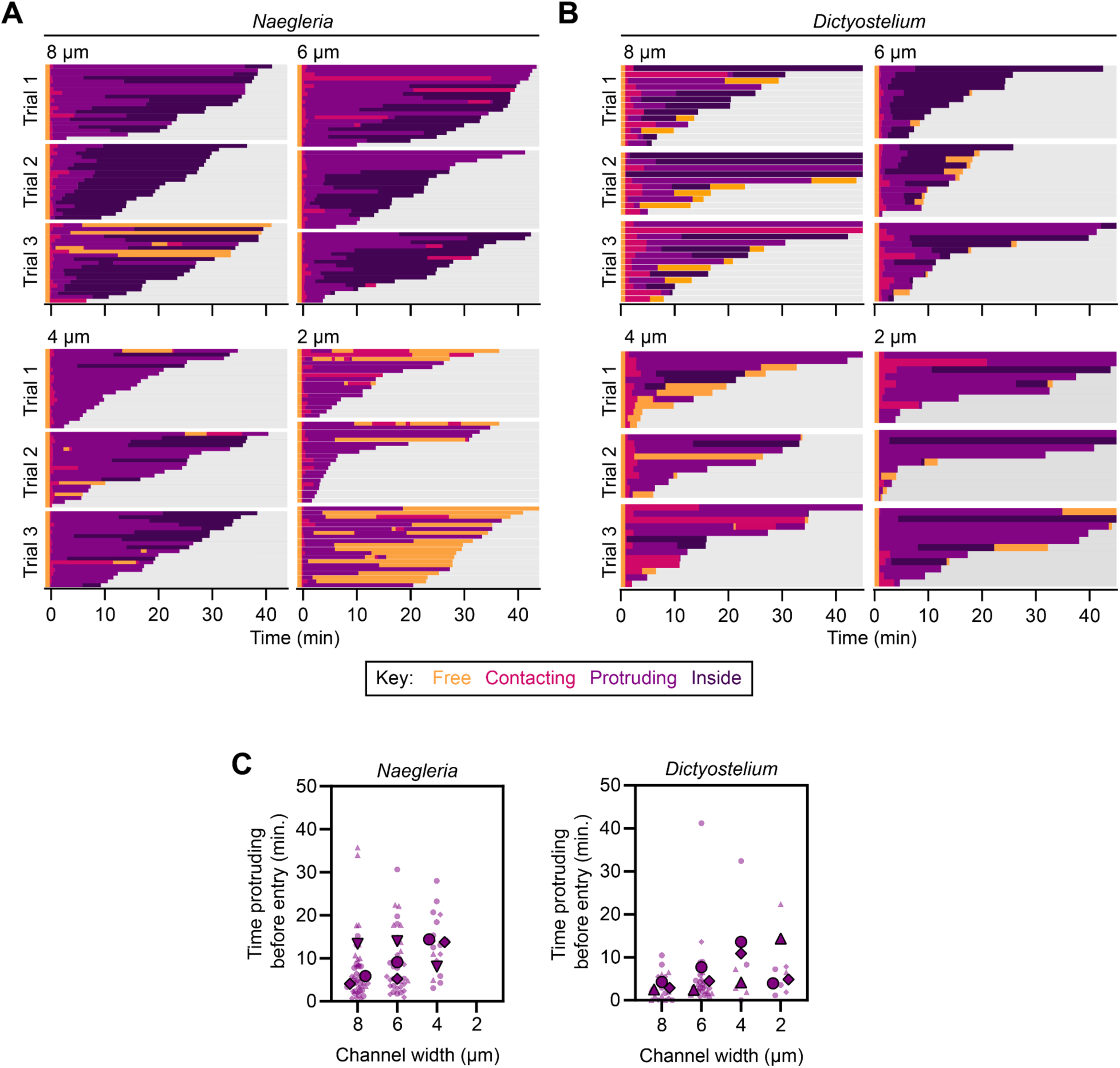
*Naegleria* enters microchannels at high rates. **(A)** *Naegleria’*s or **(B)** *Dictyostelium’*s interactions with 2-8 μm channels over 45 minutes of imaging are shown as horizontal bar plots, normalized such that t=0 represents the cell’s first contact with the interface. Trial 1 for *Naegleria* is shown in Fig. 3B. **(C)** For each cell that entered a channel, the time spent protruding prior to entry was quantified and is shown as a SuperPlot. One way ANOVAs were performed on experiment level averages, and no significant differences were detected (p>0.05).

**Figure S7.**
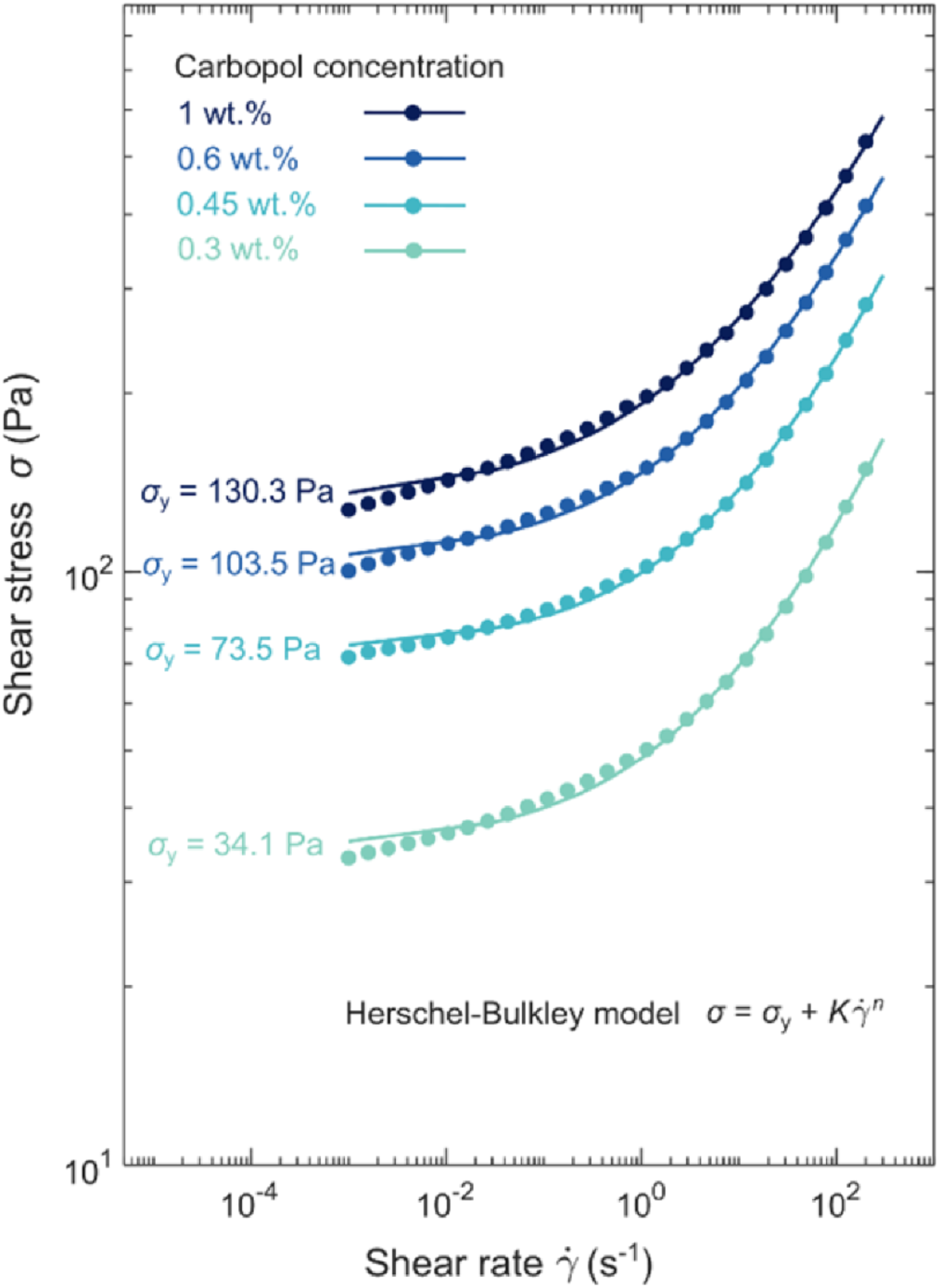
Rheological measurement of the Carbopol granular matrices. Shear stress of different Carbopol concentration versus unidirectional shear rates. The curves are fitted with the Herschel-Bulkley model for shear-thinning, viscoplastic fluids, where σy represents the yield stress, *Κ* is the consistency index, *n* is the flow index, and γ͘ is the shear rate.

## Notes

### Competing Interest Statement

The authors have declared no competing interest.

### Summary of Updates

Edits throughout the manuscript to clarify the original findings

